# A new tool CovReport generates easy-to-understand sequencing coverage summary for diagnostic reports

**DOI:** 10.1101/671511

**Authors:** Mark Gorokhov, Mathieu Cerino, Marc Bartoli, Martin Krahn, Svetlana Gorokhova

## Abstract

In order to properly interpret the results of a diagnostic gene panel sequencing test, gene coverage needs to be taken into consideration. If coverage is too low, an additional re-sequencing test is needed to make sure that a pathogenic variant is not missed. To facilitate the interpretation of coverage data, we designed CovReport, a novel easy-to-use visualization tool. CovReport generates a concise coverage summary that allows one-glance assessment of the sequencing test performance. Both gene-level and exon-level coverage can be immediately appreciated and taken into consideration for further medical decisions. CovReport does not require complex installation and can thus be easily implemented in any diagnostic laboratory setting. A user-friendly interface generates a graphic summary of coverage that can be directly included in the diagnostic report. In addition to a stand-alone version, we also provide a command line version of CovReport that can be integrated into any bioinformatics pipeline. This flexible tool is now part of routine sequencing analysis at the Department of Medical Genetics at La Timone Hospital (Marseille, France).

**Availability and implementation:** CovReport is available at http://jdotsoft.com/CovReport.php. It is implemented in Java and supported on Windows, Mac OS X and Linux.

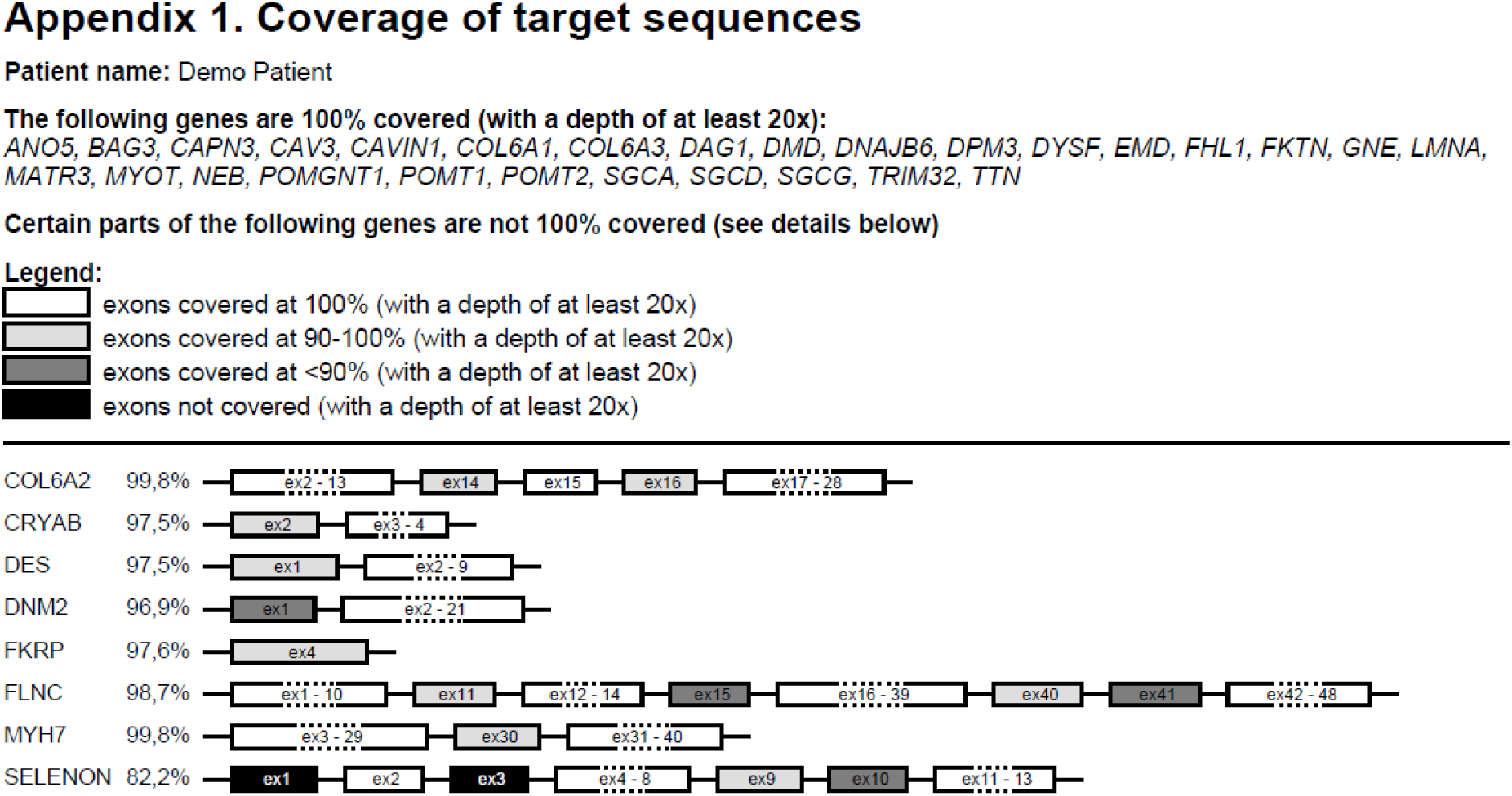

## Background

Since arrival of short-read sequencing technologies, significant efforts have been spent to improve the quality of the obtained data in order to satisfy the requirements of molecular diagnostics. However, many regions of human genome remain difficult to analyze using standard short-read sequencing approaches. These “dark” genome regions contain a number of genes responsible for human diseases [1]. For example, several neuromuscular disease-causing genes, such as *NEBULIN (NEB)* and *SELENON (SEPN1)*, overlap these difficult to sequence regions. Disease-causing variants in gene regions with suboptimal sequence coverage can be overlooked. Thus, when short-read sequencing is performed in a diagnostic setting, identifying poorly-covered regions is critical for the interpretation of a diagnostic sequencing result. If coverage is not sufficient even for a small region of a highly suspected candidate gene, an additional re-sequencing test is needed to make sure that the pathogenic variant is not missed. To facilitate the coverage data interpretation by test prescribers, we designed a novel easy-to-use visualization tool, CovReport. The concise coverage data summary generated by CovReport allows one-glance assessment of the sequencing test performance.

## Methods

### Implementation

CovReport is implemented as a standalone Java application. It can run on any platform where Java Runtime (JRE) is installed (Windows, Mac OS and Linux). The supported Java version is 8. The application is using open source external libraries (JARs) which are embedded into the main executable JAR. The following external dependencies are used: Apache PdfBox (https://pdfbox.apache.org), Apache CLI (http://commons.apache.org/proper/commons-cli), JarClassLoader (http://www.jdotsoft.com/JarClassLoader.php). The application could be downloaded from http://www.jdotsoft.com/CovReport.php as a compressed CovReport.zip file. After extraction into the local drive, the folder contains the following:

- `msg` – folder with internalization message files used for the generated PDF file
- `CovReport.jar` – Java executable archive
- `run.cmd` – Windows command to start the application
- `runFromCommandLine.cmd` – Windows command line helper

After `CovReport` first execution the following items are created:

- `pdf-results` – folder with PDF files generated by the application
- `CovReport.config` – file with configuration persistence data; this file could be manually updated and reused to replace default for command line execution

The application can be started with user interface (UI) or from command line. The command line options are:

`-i,--input <arg>` input file

`-n,--name <arg>` patient name

`-c,--config <arg>` config file (optional)

The input file containing per-exon coverage information is a tab delimited CSV file. All entries in the file can be quoted per CSV file standard or can be without quotes (there should be no special characters or in the input file). The following columns are expected: RefSeqName, GeneSymbol, Exon, Size, Mean Depth, SD Depth, Coverage 1x, Coverage 5x, Coverage 10x, Coverage 20x, Coverage 30x (see http://www.jdotsoft.com/CovReport.php for the input file example). The input file can be easily generated by the Coverage Module of VarAFT tool ([2], https://varaft.eu/) that uses Bedtools [3] to calculate exon-level coverage.

The application user interface is in English, but the generated PDF file is in the current locale language. English and French are supported in the current distribution, but other languages can be added by the user. The text included in the output pdf report can also be customized, making it possible to adopt CovReport in any diagnostic setting.

Detailed instructions for downloading and running CovReport are described at http://www.jdotsoft.com/CovReport.php

### Exome sequencing of NA12878

DNA for reference sample NA12878 was obtained from the Coriell Institute for Medical Research Repository (Coriell Institute, Camden, NJ, USA). Whole Exome Sequencing (WES) was performed by the Genomics and Bioinformatics Platform (GBiM) from the U 1251/Marseille Medical Genetics facility, using the NimbleGen SeqCap EZ MedExome kit (total design size 47 Mb) according to the manufacturer’s protocol (Roche Sequencing Solutions, Madison, USA). The SeqCap EZ MedExome kit targets the entire human exome with enhanced coverage of exons from medically relevant genes in Mendelian diseases. Enriched fragment libraries were sequenced on the Illumina NextSeq 500 platform (Illumina, San Diego, CA, USA) using a 150 bp paired-end sequencing protocol. Raw data were mapped to the built of the human genome (hg19) by using BWA 0.7.5.

## Results

CovReport allows one-glance overview of exon coverage for a diagnostic sequencing test by generating a concise easy-to-understand report in the PDF format. Intuitive interface and absence of complex installation steps makes CovReport easy to apply by users without special bioinformatics training. CovReport can also be launched by command line allowing integration into any bioinformatics pipeline. Several features make CovReport especially useful in a diagnostic setting. First, the application runs on a local computer allowing complete data security. Second, the patient’s name from the previous analysis is automatically reset upon entering the new coverage file in order to avoid identity errors. Third, the format, content and the language of the report can be easily customized by the laboratory, adapting to any diagnostic setting. Fourth, information about additional Sanger re-sequencing of suboptimally covered regions can be integrated into the coverage report of the initial short-read sequencing allowing easy tracing of sequencing experiments.

Exon coverage for genes on the panel is visualized by drawing exons shaded according to the level of coverage: 100% covered exons are white, 90-100% exons are light gray, <90% covered exons are dark gray, non-covered exons are black. Several visualization options allow adapting the graphical presentation to user’s needs. Genes with 100% coverage can be listed at the top of the report without drawing the exons (*Skip white genes* option, default). Schematic gene/exon structures will be drawn for the remaining genes, shading the lower-coverage exons. If *Merge white exons* option is selected, CovReport will fuse exons covered at 100% for more compact representation, which is useful for genes with numerous exons. Unchecking this option will produce the report with each individual exon drawn. Similarly, exons with the same shading can be merged for more compact representation (*Merge non-white exons* option). Average per-gene coverage can be shown next to each gene (*Show gene weighted coverage* option), which is also used for statistics of the total gene panel coverage (*Show statistics* option). RefSeq transcript information can be included in the report (*Show gene transcripts* option), recommended if transcripts differ in exon number leading to differences in coverage between isoforms. The default 20x depth of coverage in the report can be changed to 1x, 5x, 10x or 30x. Finally, additional comments can be included in the report.

## Use case

CovReport has been used for all diagnostic gene panel sequencing tests at the Laboratory of Molecular Genetics at the Timone Hospital (Marseille, France) since June 2018. The reports generated are directly annexed to the diagnostic results of various sequencing panels. Moreover, the coverage reports produced by CovReport are also routinely used for technical validation of the test since gene coverage statistics for the panel are integrated in the output.

To demonstrate the performance of CovReport, we used exome sequencing data for NA12878 to visualize the coverage of 40 genes on the extended Limb-Girdle Muscular Dystrophy (LGMD) gene list defined by the French National Network for Rare Neuromuscular Diseases [4]. Coverage module of VarAFT [2] was used to obtain exon coverage for these genes. The default options were used to generate the pdf report using CovReport with *Show gene transcript* and *Show statistics* options activated. As seen from Figure 1, coverage was visualized for 43 transcripts corresponding to 40 genes. Most of the genes were covered at 100%. The advantage of using *Show gene transcript* option is clear from this example, since *TTN (NM_133379)* is covered at 100%, while *TTN (NM_001267550)* has several exons with lower coverage. Indeed, *NM_133379* isoform is much shorter, as exons 50-219 of *NM_001267550* are not included in this transcript. Six exons had no regions with above 20x threshold coverage (black exons). Several other exons had suboptimal coverage in regions corresponding to more than 10% of their length (dark gray). If similar coverage of these genes is obtained after a diagnostic sequencing for a patient affected with LGMD, the prescribing clinician will take into account the presentation of the disease in order to decide if the phenotype of the patient could be potentially explained by pathogenic variants in the exons with low coverage. If that is the case, additional resequencing of these regions will be necessary.

**Figure 1.**
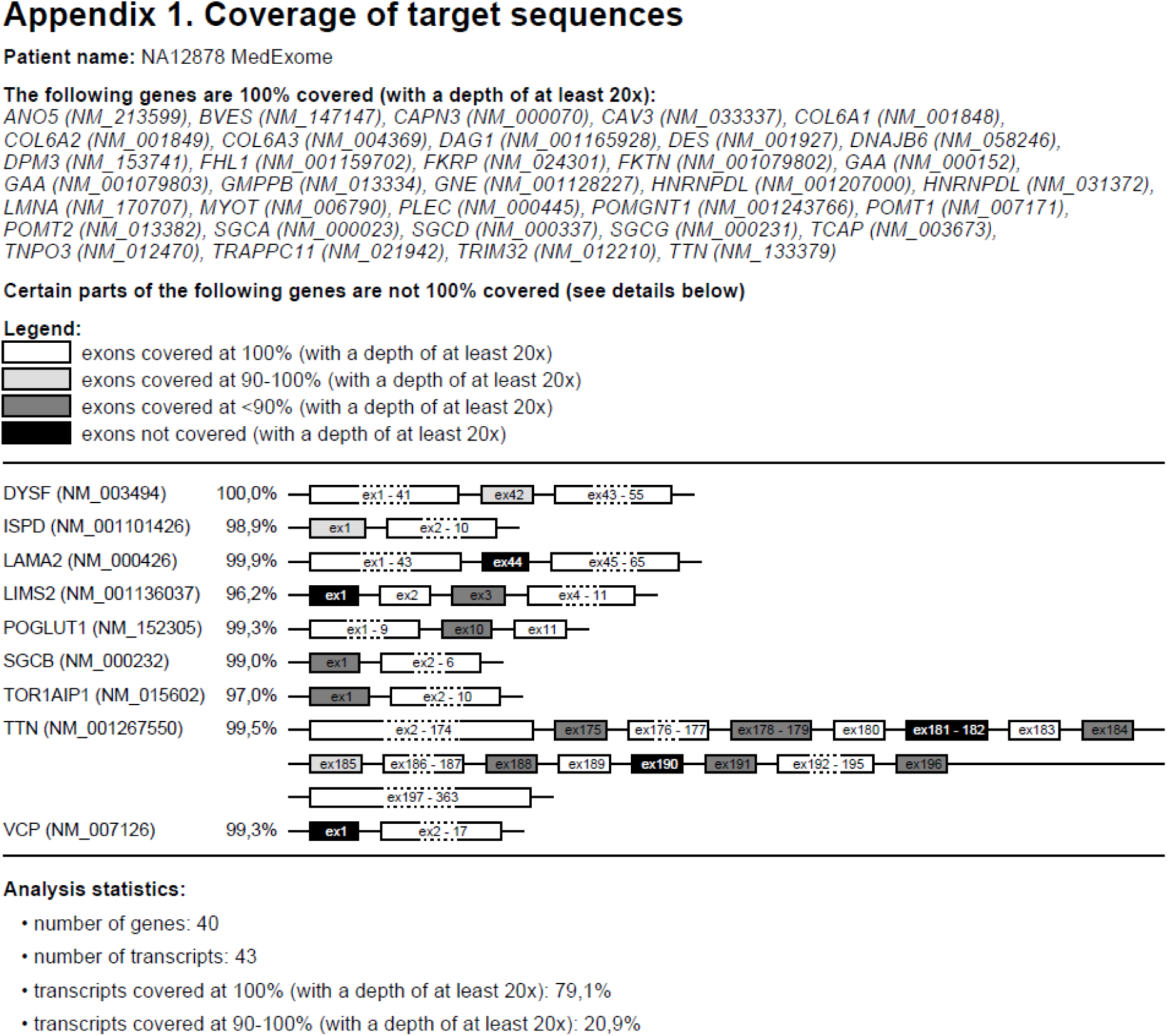
Example of a coverage report provided by CovReport.

## Discussion

Importance of coverage data is well established in genetic diagnostics. Indeed, most sequencing reports provide some information about coverage of target regions. However, these data are usually presented as an overall gene panel coverage or as an average-per-gene coverage. To our knowledge, no exon-level coverage data are provided with diagnostic reports. However, the distribution of pathogenic variants in a given gene is not homogeneous, as the disease-causing amino acid changes are often clustered in functionally important regions of the protein. Thus, information about coverage of the regions where disease-causing variants are concentrated is much more informative than an overall average gene coverage [5]. CovReport highlights individual exons with suboptimal coverage, facilitating the interpretation of the sequencing data.

The ultimate goal of diagnostic sequencing is to obtain an above-threshold coverage for the entire genomic area targeted by a given gene panel. This is feasible for small panels with genes that can be robustly sequenced. However, this goal is much harder to achieve for larger gene panels containing difficult to sequence genes. This is the case for the neuromuscular disorder field, since 33 out of 203 genes on the consensus myopathy gene lists [4] contain “dark” regions of the genome that are not easily accessible using standard short-read sequencing approaches [1]. Disease-causing variants in these regions can therefore be overlooked leading to a false negative molecular diagnostic result. CovReport effectively highlights and draws attention to the exons with suboptimal coverage, allowing the prescriber to evaluate the need for a complementary re-sequencing test for these areas.

Given the critical importance of coverage data, several tools have been previously developed to evaluate the coverage of target regions after short-read sequencing in a diagnostic setting [2,6,7]. While these tools are useful to monitor quality control of runs and samples during technical validation of sequencing tests, unlike CovReport, they do not provide exon-level coverage visualization that can be directly annexed to the diagnostic report.

In conclusion, CovReport generates a one-glance graphical overview of coverage for individual exons on a gene panel, facilitating interpretation of the sequencing test. CovReport is flexible and easy-to-use, making it easy adopt in any diagnostic setting.

## Acknowledgements

We would like to thank the initial users of CovReport at the Laboratory of Molecular Genetics at the Timone Hospital, Marseille (Caroline Lacoste, Véronique Blanc, Christope Pecheux and Amandine Boyer) as well as Henri Pégeot at the Laboratory of Molecular Genetics, Montpellier, France for the valuable comments and suggestions for improvement. We would also like to thank the Genomics&Bioinformatics Platform at MMG for providing the exome sequencing data for NA12878.

## Notes

http://www.jdotsoft.com/CovReport.php

